# Reovirus efficiently reassorts genome segments during coinfection and superinfection

**DOI:** 10.1101/2022.06.09.495584

**Authors:** Timothy W. Thoner, Madeline M. Meloy, Jacob M. Long, Julia R. Diller, James C. Slaughter, Kristen M. Ogden

## Abstract

Reassortment, or genome segment exchange, increases diversity among viruses with segmented genomes. Previous studies on the limitations of reassortment have largely focused on parental incompatibilities that restrict generation of viable progeny. However, less is known about whether factors intrinsic to virus replication influence reassortment. Mammalian orthoreovirus (reovirus) encapsidates a segmented, double-stranded RNA genome, replicates within cytoplasmic factories, and is susceptible to host antiviral responses. We sought to elucidate the influence of infection multiplicity, timing, and compartmentalized replication on reovirus reassortment in the absence of parental incompatibilities. We used an established post-PCR genotyping method to quantify reassortment frequency between wild-type and genetically-barcoded type 3 reoviruses. Consistent with published findings, we found that reassortment increased with infection multiplicity until reaching a peak of efficient genome segment exchange during simultaneous coinfection. However, reassortment frequency exhibited a substantial decease with increasing time to superinfection, which strongly correlated with viral transcript abundance. We hypothesized that physical sequestration of viral transcripts within distinct virus factories or superinfection exclusion also could influence reassortment frequency during superinfection. Imaging revealed that transcripts from both wild-type and barcoded viruses frequently co-occupied factories with superinfection time delays up to 16 hours. Additionally, primary infection dampened superinfecting virus transcription with a 24 hour, but not shorter, time delay to superinfection. Thus, in the absence of parental incompatibilities and with short times to superinfection, reovirus reassortment proceeds efficiently and is largely unaffected by compartmentalization of replication and superinfection exclusion. However, reassortment may be limited by superinfection exclusion with greater time delays to superinfection.

**IMPORTANCE:** Reassortment, or genome segment exchange between viruses, can generate novel virus genotypes and pandemic virus strains. For viruses to reassort their genome segments, they must replicate within the same physical space by coinfecting the same host cell. Even after entry into the host cell, many viruses with segmented genomes synthesize new virus transcripts and assemble and package their genomes within cytoplasmic replication compartments. Additionally, some viruses can interfere with subsequent infection of the same host or cell. However, spatial and temporal influences on reassortment are only beginning to be explored. We found that infection multiplicity and transcript abundance are important drivers of reassortment during coinfection and superinfection, respectively, for reovirus, which has a segmented, double-stranded RNA genome. We also provide evidence that compartmentalization of transcription and packaging is unlikely to influence reassortment, but the length of time between primary and subsequent reovirus infection can alter reassortment frequency.

## INTRODUCTION

Genome segment reassortment is a major mechanism of genetic diversity acquisition among viruses with segmented genomes. Reassortment events can engender the generation of pandemic virus strains (1, 2), provide avenues for zoonotic transmission events (reviewed in (3)), and increase the prevalence of antiviral drug-resistant variants (4, 5). Reassortment has been observed for viruses belonging to nearly every family with segmented genomes, including the *Arenaviridae* (6, 7), *Bunyaviridae* ((8), reviewed in (9)), *Orthomyxoviridae* (reviewed in (10)), and *Reoviridae* (11–15) families. While reassortment occurs often in nature, limitations to gene segment exchange do exist. Specifically, it is well-established that reassortment events can disrupt critical interactions between viral RNA and proteins, leading to virus progeny that are less fit than parental viruses or are completely nonviable (16–19); reviewed in (20)). Limitations to reassortment as a result of parental incompatibilities are referred to as “segment mismatch”. While the influence of segment mismatch on reassortment is well-documented, other factors, which we refer to as “intrinsic influences,” may also contribute to the efficiency of reassortment even after a cell has been coinfected. Less is understood about intrinsic influences of reassortment, though some mechanisms, such as physical separation of viruses within coinfected cells, have been proposed (21).

Reovirus is a member of the *Reoviridae* family and encapsidates a genome composed of ten double-stranded RNA segments named according to their respective sizes – large (L1, L2, L3), medium, (M1, M2, M3), and small (S1, S2, S3, S4). Reovirus reassortment has previously been described as a nonrandom process, such that the frequency of reassortment events is lower than expected, and specific gene constellations are preferred in coinfection progeny (19, 22). Thus, reovirus reassortment appears to be constrained by segment mismatch.

Many RNA viruses compartmentalize replication processes within membranous and proteinaceous bodies in the cytoplasm, which may protect from antiviral responses and concentrate factors important for efficient viral replication. Reovirus replication occurs in cytoplasmic virus factories (VFs), which are assembled from interactions between non-structural proteins µNS and σNS and act as the primary site of viral positive-sense RNA (+RNA) synthesis, genome packaging, and new particle assembly (23, 24). Recent studies indicate that reovirus +RNA localizes to both the cytoplasm and VFs and that non-structural protein σNS is responsible for recruiting +RNA to VFs (25). Reovirus VFs are not static. First appearing as small, punctate bodies in the cytoplasm, VFs fuse and become larger as replication progresses (26, 27). However, it is unclear if +RNA traffics out of and between VFs and what effect trafficking may have on the capacity of transcripts from coinfecting viruses to colocalize and be copackaged into assembling virions. Thus, VFs may influence reassortment frequency either by sequestering viral RNA to prevent reassortment events or by facilitating the accumulation of viral RNA from coinfecting viruses to promote reassortment events. Recent work suggests that VF morphology is not an important determinant of reassortment frequency during simultaneous coinfection (28). However, it is unknown whether newly synthesized viral RNA can enter mature VFs and, thus, how VFs affect reassortment during superinfection, when the timing of coinfection is asynchronous.

Superinfection exclusion, also known as viral interference, may also influence reassortment frequency when coinfection does not occur simultaneously. Superinfection exclusion occurs when infection with a first virus interferes with subsequent infection of the same cell or organism by a second virus. Multiple mechanisms of superinfection exclusion have been identified, including competition for host resources (29), degradation or downregulation of entry receptors (17, 30, 31), and antiviral host responses (32, 33). Previous studies of reovirus reassortment have suggested that superinfection is not excluded, as reassortant progeny can be detected even when there is a substantial time delay separating primary infection and superinfection (34). However, type 3 reovirus is known to potently induce and to be susceptible to the antiviral effects of type 1 and type 3 interferons (35–37). Whether reovirus limits superinfection, and what affect this may have on reassortment, is an open question.

In the current study, we sought to determine whether processes intrinsic to the reovirus replication cycle influence reassortment by examining reovirus reassortment in the absence of segment mismatch. We used a post-PCR genotyping method to quantify reassortment following coinfection and superinfection of cultured cells with type 3 reoviruses in wild-type and genetically-barcoded forms. We also determined viral RNA localization relative to VFs and quantified viral RNA abundance during coinfection and superinfection. Consistent with published data, we found that reassortment events are frequent during high multiplicity coinfection in the absence of segment mismatch (28).

However, reassortment frequency decreased as the time delay to superinfection was increased. During superinfection, the time to introduction of the superinfecting virus and the abundance of superinfecting virus +RNA displayed strong positive correlations with reassortment frequency. Furthermore, +RNA imaging during coinfection and superinfection revealed pools of cytoplasmic and VF-localized reovirus transcripts, and VFs did not appear to pose a significant barrier to reassortment events. Superinfection exclusion was not detected when primary infection and superinfection occurred within a single replication cycle. However, with greater time delay to superinfection, type 3 reovirus primary infection did reduce the abundance of superinfecting reovirus transcripts. These findings suggest that infection multiplicity is a key determinant of reassortment during synchronous coinfection. Further, compartmentalization of replication is not a critical mediator of reassortment potential following superinfection; rather, the abundance of superinfecting virus transcripts appears to dictate reassortment frequency during superinfection. Lastly, superinfection exclusion is unlikely to influence reassortment during a single replication cycle but may influence reassortment potential in certain contexts.

## MATERIALS AND METHODS

### Cells and Antibodies

Spinner-adapted murine L929 fibroblasts (L cells) were grown in suspension culture in Joklik’s minimum essential medium (JMEM) (US Biological) supplemented to contain 5% fetal bovine serum (FBS) (Gibco) and 2 mM L-glutamine. Baby hamster kidney cells expressing T7 RNA polymerase under control of a cytomegalovirus promoter (BHK-T7) (38) were grown in Dulbecco’s modified Eagle’s minimal essential medium (Corning) supplemented to contain 5% FBS and 2 mM L-glutamine. These cells were propagated in the presence of 1 mg/ml G418 (Invitrogen) during alternate passages.

Rabbit polyclonal reovirus antiserum and rabbit σNS-specific antiserum (39) were gifts from Dr. Terence Dermody.

### Viruses

Reovirus strain rsT3D^I^ is a variant of human reovirus laboratory strain rsT3D in which a T249I mutation has been introduced into the attachment protein, σ1, rendering it resistant to trypsin proteolysis (40). A plasmid encoding T3D S1 in which a T249I mutation had been introduced into σ1 was engineered from the parental reverse genetics plasmid using ‘round the horn PCR (https://openwetware.org/wiki/%27Round-the-horn_site-directed_mutagenesis) with mutagenic primers (sequences available upon request) and Phusion High-Fidelity DNA Polymerase (New England Biolabs). Plasmids encoding rsT3D^I^ segments with silent barcode mutations (Table 1) were engineered from the parental rsT3D^I^ reverse genetics plasmids and pBac T7 rsT3D S1 T249I using ‘round the horn PCR with mutagenic primers (sequences available upon request) and Phusion High-Fidelity DNA Polymerase. Reovirus strains rsT3D^I^ (WT) and genetically barcoded rsT3D^I^ BC (BC) were prepared using plasmid-based reverse genetics (38, 40). Briefly, plasmids encoding all ten reovirus genome segments were transfected into BHK-T7 cells using TransIT-LT1 (Mirus Bio). After the first signs of cytopathic effects, virus was released from cells by multiple rounds of freezing at -80°C and thawing at room temperature. Individual clones were then isolated by plaque assay. At least three clones of each recombinant virus strain were isolated by plaque purification and propagated in L cells for two passages as described (41). Viral titers were determined by plaque assay using L cells (41). Virus particles were purified from infected L cells by Vertrel XF (DuPont) extraction and CsCl gradient centrifugation (modified from (41). The presence of engineered mutations was confirmed by Sanger sequencing following extraction of RNA from virus stocks with TRIzol (Invitrogen) and cDNA amplification using a OneStep RT-PCR Kit (Qiagen) and segment-specific primers (sequences available upon request), according to manufacturer protocols.

**Table 1.**
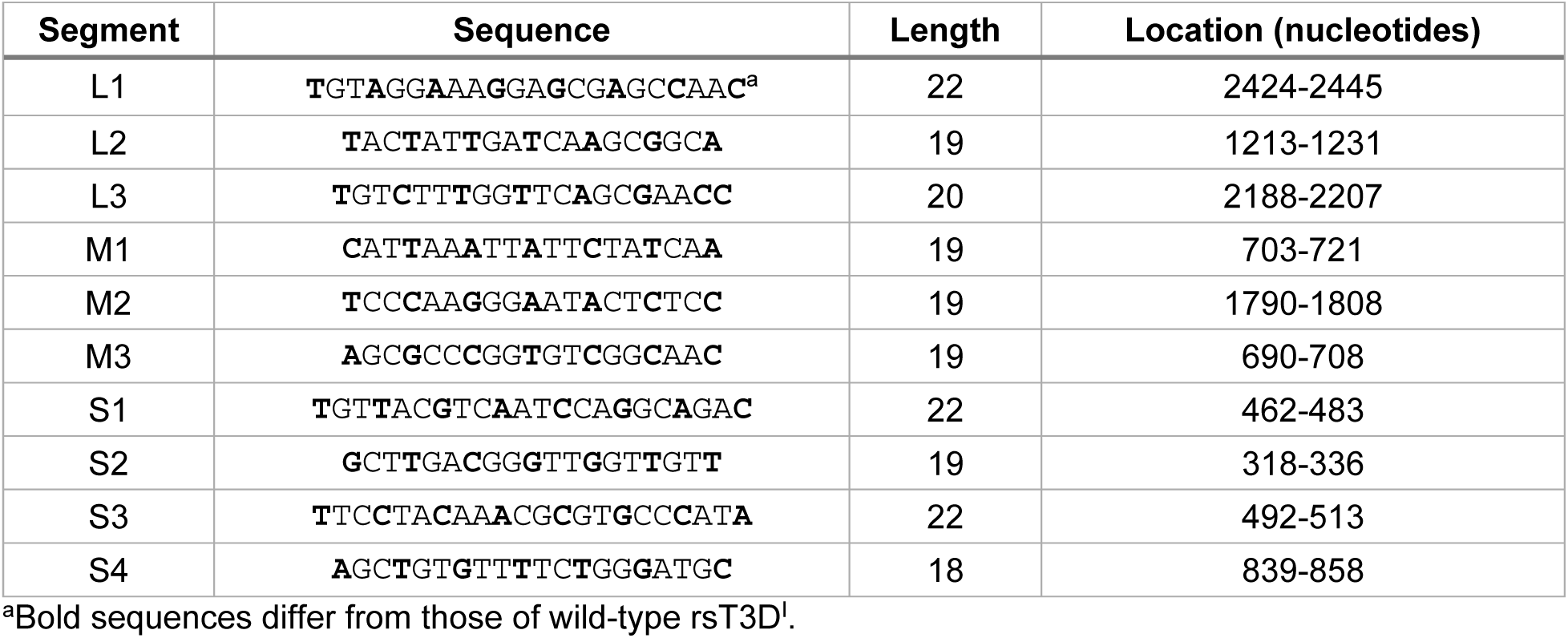
Barcode sequences and locations.

### Multi-step replication

L cells (8 × 10^5^ cells per well) in 12-well plates were adsorbed with stocks generated from three independent clones each of WT and BC diluted in phosphate-buffered saline (PBS) to a MOI of 0.1 plaque-forming units (PFU) per cell for 1 h at room temperature. Inocula were aspirated, cells were washed, and 1 ml of JMEM supplemented to contain 5% FBS (complete JMEM) per well was added. Cells were incubated for 0, 8, 12, 16, 20, 24, or 48 h prior to two cycles of freezing at -80°C and thawing at room temperature, to lyse cells. Virus titer in cell lysates was determined by plaque assay. Two independent experiments were conducted. Data at each time point were compared by unpaired student’s *t* test.

### Coinfection experiments

L cells (4 × 10^5^ cells per well) in 12-well plates were adsorbed with WT, BC, or both viruses. Virus was diluted in complete JMEM to a MOI of 0.1, 0.5, 1, 10, or 100 PFU per cell per virus for 1 h at room temperature. Cells were washed, and 1 ml per well of complete JMEM was added. Leftover inocula were stored at 4°C for back-titration. Cells were incubated for 24 h prior to two cycles of freezing at - 80°C and thawing at room temperature to lyse cells. Individual viral progeny were isolated by plaque purification (41). Plaques were vortexed in 0.25 ml complete JMEM and amplified by adsorption on L cell monolayers in 12-well plates. Total RNA was extracted from monolayers at 48-72 h post infection (p.i.) using TRIzol (Invitrogen), according to the manufacturer protocol. Viral +RNA was quantified by RT-qPCR, and parental segment origins were determined by high-resolution melt (HRM) analysis, as described below.

### Superinfection experiments

L cells (4 × 10^5^ cells per well) in 12-well plates were adsorbed with WT diluted in complete JMEM to a MOI of 10 PFU per cell for 1 h at room temperature. Cells were washed with 1 mL complete JMEM, and at 0, 4, 8, 12, or 16 h p.i., the cells were superinfected with BC diluted in complete JMEM to a MOI of 10 PFU per cell. Cells were washed, and 1 ml per well of complete JMEM was added. Leftover inocula were stored 4°C for back-titration. At 24 h post primary infection, two cycles of freezing at -80°C and thawing at room temperature were conducted to lyse cells.

Individual viral progeny were isolated by plaque purification (41). Plaques were vortexed in 0.25 ml complete JMEM and amplified by adsorption on L cell monolayers in 12-well plates. Total RNA was extracted from monolayers at 48-72 h p.i. using TRIzol (Invitrogen), following manufacturer’s protocol. Viral +RNA was quantified by RT-qPCR, and parental segment origins were determined by HRM analysis, as described below.

### Viral genotyping by high resolution melt analysis

RNA extracted from a single amplified plaque was randomly primed, and cDNA was generated using SuperScript III reverse transcriptase (Invitrogen). cDNA was amplified using MeltDoctor HRM Master Mix (Applied Biosystems) and primers surrounding either the wild-type or barcoded region of the reovirus S1 (F 5’-CATTGACCACCGAGCTATC-3’; R 5’- CCCATGGTCATACGGTTATT-3’), S2 (F 5’-ATTCCGTTCCGTCCTAAC-3’; R 5’- CAGTCGTACACTCGATCTG-3’), S3 (F 5’-CACTTGCCAGATTGTTTACC-3’; R 5’- CAGCGTCATACAGTCCAA-3’), M1 (F 5’-GTGTTCCTCACTCTCGATTT-3’; R 5’- GCAGACGCTTTCTGTTTATC-3’), M3 (F 5’-TCTGACGCTAAAGGGATAATG-3’; R 5’- GCAGTCTCCAAGGTGAAATA-3’), or L2 (F 5’-AGACTCGGCATGAGAATATC-3’; R 5’-TAACGGGAGTTGCGTAAG -3’) genome segments. Melt curves were generated following cDNA amplification and WT and BC genotyping was performed with High Resolution Melt Software ver. 3.0.1 (Applied Biosystems). All clones were run in duplicate or triplicate to validate the called genotype. Most clones were consistently called as either wild-type or barcode for a given segment. Clones that consistently melted at a temperature in between the expected melt temperatures for the wild-type and barcode genotypes, yielding an ambiguous parental genotype, were removed from the analysis. Such clones represented a small fraction (2.8%) of total isolated plaques. Studies of genetically-marked polioviruses revealed that virion aggregates led to 5-7% of total plaques being formed by more than a single virion (42). Thus, clones with an ambiguous parental origin may represent plaques that were formed by aggregated WT and BC virions. Plaques amplified from cells infected with only WT or BC were included in each run, in triplicate, as genotype references.

### Quantification of coinfectivity by branched DNA FISH and fluorescent focus assay

L cells (2 x 10^4^ cells per well) in 96-well black-walled plates were adsorbed with 0.1, 0.5, 1, 10, or 100 PFU WT and BC reovirus for 1 hour at room temperature. Cells were washed with complete JMEM, and infection was allowed to proceed for 24 h.

Infected cells were fixed and stained using branched DNA FISH probes specific to the wild-type or barcoded region of the T3D reovirus L1, S3, and S4 +RNA, per manufacturer’s protocol. Cells were then stained with DAPI in PBS for 10 minutes at room temperature before washing with PBS and imaging on the ImageXpress high-content imaging system (Molecular Devices). Coinfectivity was determined using a custom module in MetaXpress software by thresholding images and quantifying and averaging the percentage of cells from four fields of view positive for AlexaFluor488 (WT T3D^I^) and AlexaFluor647 (BC T3D^I^).

### Quantitative reverse transcription PCR (RT-qPCR)

L cell monolayers were disrupted by scraping and pelleted at 200 x g for 5 min before RNA was isolated from 2 x 10^5^ cells using the RNeasy Plus Mini RNA extraction kit (Qiagen) per manufacturer’s protocol.

RNA concentration was quantified by Nanodrop. Equal amounts of total RNA (400 ng) were primed with random hexamers (Invitrogen), and cDNA was generated using SuperScript III reverse transcriptase (ThermoFisher) using manufacturer’s protocol. For viral RNA standards, RNA was extracted from WT and BC T3D^I^ viruses purified by cesium-chloride gradient ultracentrifugation, normalized based on concentration, serially diluted to achieve target standard curve concentrations, and reverse transcribed. cDNA was amplified and quantified using PowerUP SYBR Green Master Mix (ThermoFisher) and primers specific to the wild-type or barcoded region of the reovirus S4 gene (WT F: 5’-GGCCGTATTCTCAGGAATGTT-3’; BC F: 5’-AGCTGTGTTTTCTGGGATGC-3’; WT/BC R: 5’-AATCTTCTCGACACCCCAAG-3’). To calculate the concentration of RNA, the concentration of total RNA from standard curve stocks was quantified by Nanodrop. Genome segments are anticipated to be present in equimolar concentrations in purified particle preparations (43). The concentration of S4 was calculated as a proportion of total viral genome length. C_T_ values of serially diluted standards were determined, and the concentrations of all other samples were interpolated relative to standards of known concentration.

### Branched DNA FISH staining and image analysis of +RNA and VFs

L cells were seeded onto coverslips in a 24-well plate and infected with MOI = 10 PFU per cell per virus of WT and BC at indicated time points. Infected cells were fixed and stained to detect S3, S4, and L1 +RNAs from WT and BC, and viral nonstructural protein σNS to identify VFs, using the ViewRNA Cell Plus assay kit (ThermoFisher) according to manufacturer’s recommendations. Coverslips were mounted using Prolong Gold antifade reagent with DAPI (Invitrogen). Confocal imaging was performed with a Zeiss LSM880 confocal microscope equipped with 40× 1.30 C Plan-Apochromat Oil objective lens. Image analysis was conducted with FIJI (v.1.53c). VFs were identified by performing a gaussian blur and subtracting background to remove low intensity signal in the cytoplasm associated with σNS, thresholding on the σNS channel, and analyzing particles. Regions of interest corresponding to VFs were compiled for each analyzed cell. VFs identified in this manner were visually confirmed to represent true staining before determining whether VFs contained WT and BC +RNA. To quantify the proportion of +RNA within and outside of VFs, raw integrated density associated with WT and BC +RNA within an individual cell, and within all VFs within that cell, was measured. To determine whether VFs contained +RNA from WT and BC, background was subtracted, and images were thresholded on the WT +RNA (AlexaFluor488) and BC +RNA (AlexaFluor647) channels. The regions of interest identifying VFs from each cell were overlayed onto the thresholded +RNA channels and the mean fluorescence intensity of +RNA signal within VFs was quantified. VFs that contained +RNA corresponding to the AlexaFluor488 channel, AlexaFluor647 channel, or both channels were called as WT+, BC+, or WT+BC+, respectively. Data were collected for a total of 30 cells and ∼1200-1500 VFs at each time point.

### Statistical analyses

Binomial analyses were used to determine whether reassortment occurred randomly. Given that reassortment was assessed for six genome segments, and there were two possible genotypes for each segment, either WT or BC, there were 2^6^, or 64, possible combinations of segments. Two (3%) of these 64 combinations would represent the WT and BC parental genotypes, while the other 62 (97%) would represent reassortant genotypes. If genome segments were randomly exchanged, the number of reassortant progeny should not significantly deviate from the expected percentage of reassortant progeny.

To determine whether larger or small genome segments undergo reassortment more frequently, genome segments from each clone were divided into triplets based on size – small segments (S1, S2, and S3) and medium/large segments (M1, M3, and L2). The percentage of segments of WT origin for medium/large and small segments, and the percentage of medium/large and small segment triplets that were reassortant, were totaled for all coinfection and superinfection progeny. Chi squared analyses were then used to determine if the observed number of medium/large segments that were i) of WT origin, or ii) were reassortant, differed from that observed for small segments.

Comparison of small and medium/large segments revealed no statistically significant differences in the proportion of segments that were WT during coinfection at any MOI (MOI = 0.1 PFU/cell/virus, p = 0.937; MOI = 1 PFU/cell/virus, p = 0.813; MOI = 10 PFU/cell/virus, p = 0.644; MOI = 100 PFU/cell/virus, p = 0.318) or during superinfection at any time point (0 h superinfection= 0.735; 4 h superinfection= 0.537; 8 h superinfection= 0.248; 12 h superinfection= 0.572; 16 h superinfection= 0.538). Additionally, no differences were observed for the percentage of clones that were reassortant between medium/large and small segments during coinfection (MOI = 0.1 PFU/cell/virus, p = 0.343; MOI = 1 PFU/cell/virus, p = 1.0000; MOI = 10 PFU/cell/virus, p = 0.813; MOI = 100 PFU/cell/virus, p = 0.634) or superinfection (0 h superinfection= 0.623; 4 h superinfection= 0.114; 8 h superinfection= 0.095; 12 h superinfection= 0.442; 16 h superinfection= 0.343).

Simple linear regression analyses were conducted to compare transcript abundance to reassortment frequency during coinfection and superinfection. To determine if primary infection restricted superinfecting virus transcript abundance at a 24 h superinfection time point, statistical significance was determined by unpaired t-test. One-way ANOVA with Tukey’s multiple comparisons test was conducted to determine whether significant differences existed between time points for the proportion and ratio of VFs containing +RNA from WT and BC viruses. For experiments comparing transcript abundance in mock primary infection and BC primary infection over a time course, statistical significance was determined by two-way ANOVA with Sidak’s multiple comparison test. All statistical analyses were conducted using GraphPad Prism version 9.3.1.

## RESULTS

### A genetically barcoded reovirus displays identical replication kinetics and can be differentiated from wild-type reovirus during coinfection

To quantify reassortment frequency in the absence of protein and RNA incompatibilities, we engineered a barcoded rsT3D^I^ reovirus (BC) that is isogenic to wild-type rsT3D^I^ (WT) except for 7 or 8 synonymous changes within a 21-24 nucleotide region of each viral genome segment (**Table 1**). Barcodes are distal from terminal sequences required for packaging, assortment, transcription, and translation and are not anticipated to alter RNA folding or recognition in a way that will diminish viral fitness (**Fig. 1A**). To determine whether barcoding alters BC reovirus replication, we quantified virus titer from three plaque-purified clones of BC over a time course and found that these clones replicate with nearly identical kinetics to WT in murine L929 fibroblasts (L cells) (**Fig. 1B**). To differentiate WT and BC genomes in coinfected cells and quantify reassortment frequency, we used high-resolution melt analysis (HRM). HRM is a post-PCR genotyping method that enables the detection of genetic variants based on the melt temperature of PCR-amplified cDNA products and has been used to study reassortment of genetically barcoded influenza virus and reovirus (44, 45). We extracted RNA from purified WT and BC reovirus stocks and conducted HRM for each genome segment using primers that amplify a region surrounding the genetic barcode. Melt temperatures for four genome segments were too similar to consistently differentiate the WT and BC sequences. However, difference plots of melt curves showed that six WT and BC genome segments could be easily differentiated after PCR amplification (**Fig. 1C**). Thus, we developed a system useful for quantifying reovirus reassortment, as the WT and BC viruses replicate with equivalent efficiency but six of their genome segments can be consistently distinguished by HRM.

**Figure 1.**
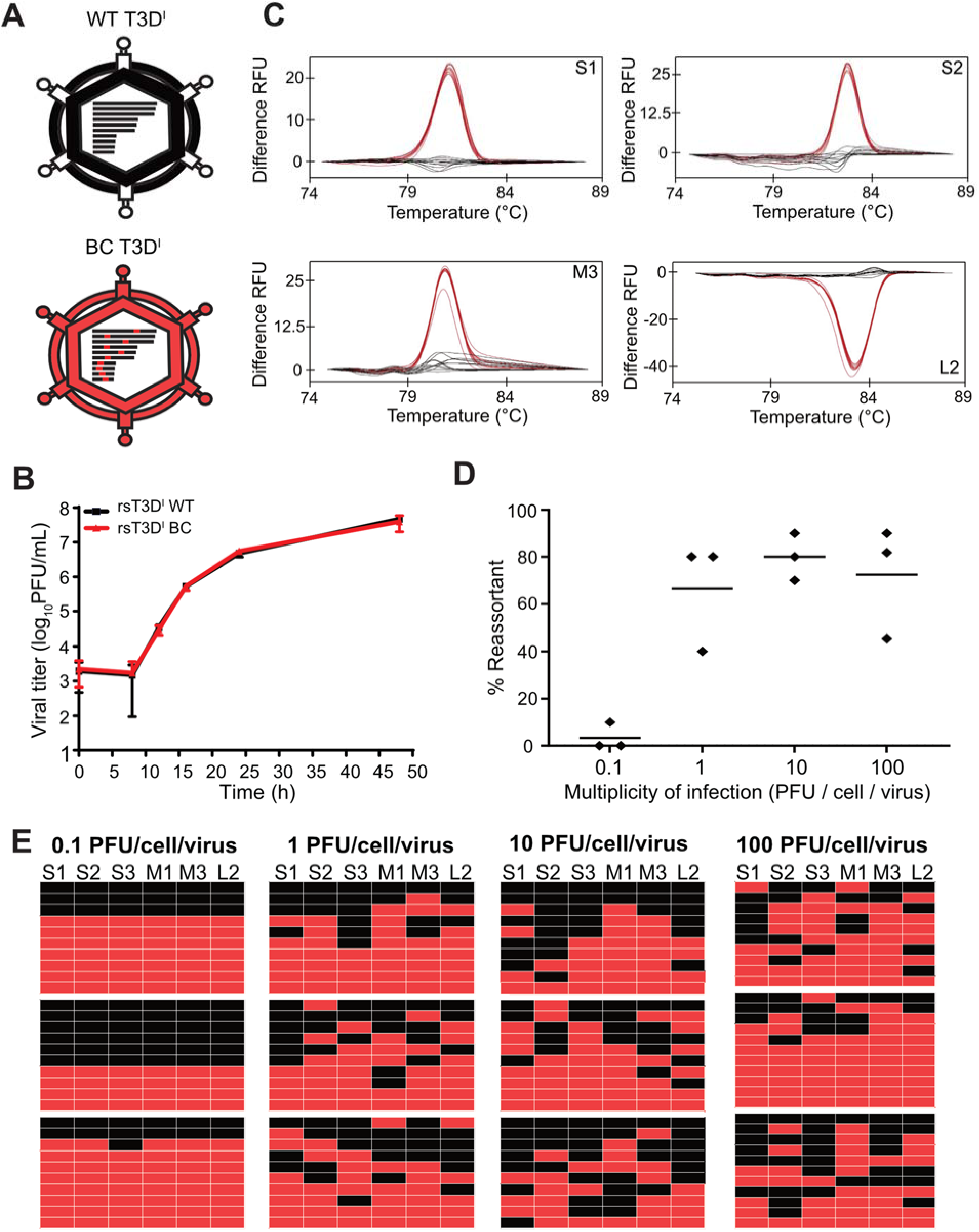
Reovirus reassortment is frequent during high multiplicity coinfection. (A) Schematic representation of reovirus virions, genome segments, and barcoding strategy. Black bars represent genome segments, with red representing barcodes, or regions into which a series of silent, single-nucleotide substitutions were introduced. (B) L cells were adsorbed with WT or BC reovirus at a multiplicity of 0.1 PFU per cell and incubated for the indicated times prior to cell lysis. Virus titer in cell lysates was determined by plaque assay. n = 3 pairs of plaque-purified clones from each of two independent experiments. (C) Representative HRM difference plots indicating fluorescence relative to a WT control over a range of temperatures for indicated genome segments. n = 4 WT or BC clones, analyzed in triplicate. Black lines indicate WT genome segment. Red lines indicate BC genome segment. (D) L cells were coinfected with WT and BC reovirus at a MOI of 0.1, 1, 10, or 100 PFU per cell per virus before quantifying reassortment frequency using HRM. (E) Honeycomb plots indicate genotypes of individual coinfection progeny viruses analyzed from the experiment in (D). Each row represents an independently isolated clone. Each column represents the indicated genome segment. Black indicates WT genome segments, and red indicates BC genome segments. n = 3 independent experiments, with at least 10 progeny analyzed per experiment.

### Reovirus reassortment occurs frequently during simultaneous coinfection

To quantify the frequency of reovirus reassortment in the absence of segment mismatch, we simultaneously coinfected L cells with WT and BC reovirus at increasing multiplicities of infection (MOI). After 24 h, infectious progeny from these coinfections were isolated by plaque assay and amplified, their RNA was extracted, and the parental origin of the S1, S2, S3, M1, M3, and L2 segments was determined by HRM. At a MOI of 0.1 PFU per cell per virus, reassortment events were rare, with only 3% of progeny clones packaging a detectably reassorted genome (**Fig. 1D-E**). However, reassortment frequency increased with higher MOI; 67%, 80%, and 72% of clones were reassortants at MOIs of 1, 10, or 100 PFU per cell per virus, respectively (**Fig. 1D-E**). Thus, segment exchange occurred frequently beginning at a MOI of 1 PFU per cell per virus, and increasing the MOI above this level did not substantially enhance reassortment frequency. Using HRM, we may not have detected every reassortant virus among our progeny, since four of the reovirus genome segments were not analyzed. However, we mathematically adjusted our expected results to account for analysis of only six of the ten reovirus genome segments. When six segments are analyzed, there are 2^6^, or 64, possible genome segment combinations. If reassortment was statistically random, 62 of every 64 virus progeny (∼97%) would package a reassortant genotype, while only two would package the parental (WT or BC) genotype. Since reassortment frequency peaked at ∼80%, in our assay it failed to reach a frequency suggestive of entirely random genome segment exchange. We were also curious whether certain genome segments are exchanged more frequently than others. Binomial and Χ^2^ analyses indicate that there was no preference for certain segment size classes (L/M or S) to reassort more frequently (**Materials and Methods**). These findings suggest that following simultaneous coinfection, reovirus reassortment occurs frequently during a single cycle of replication, though there may be some limitations to this process that are unlikely to involve viral protein or RNA incompatibilities.

### RNA abundance fails to explain reassortment frequency during coinfection

The abundance of viral RNA from each coinfecting virus within a cell may influence reassortment frequency, as progeny virus particles are more likely to package abundant viral +RNA molecules. To quantify the abundance of WT and BC RNA transcripts during coinfection, we used RT-qPCR. Specifically, we designed primers targeting either the WT sequence or the barcoded region of the reovirus S4 segment that would allow for specific amplification of RNA from each virus. Experiments in which cells were uninfected, singly infected, or coinfected with both WT and BC indicated that the primers yielded low background amplification when the target virus was absent (C_T_ >28), but substantial amplification occurred when the target virus was present (C_T_ ∼12- 16) (**Fig. 2A-B**). These findings suggested that the WT- and BC-specific primers could accurately differentiate between the two viruses during coinfection to permit estimation of viral RNA abundance.

**Figure 2.**
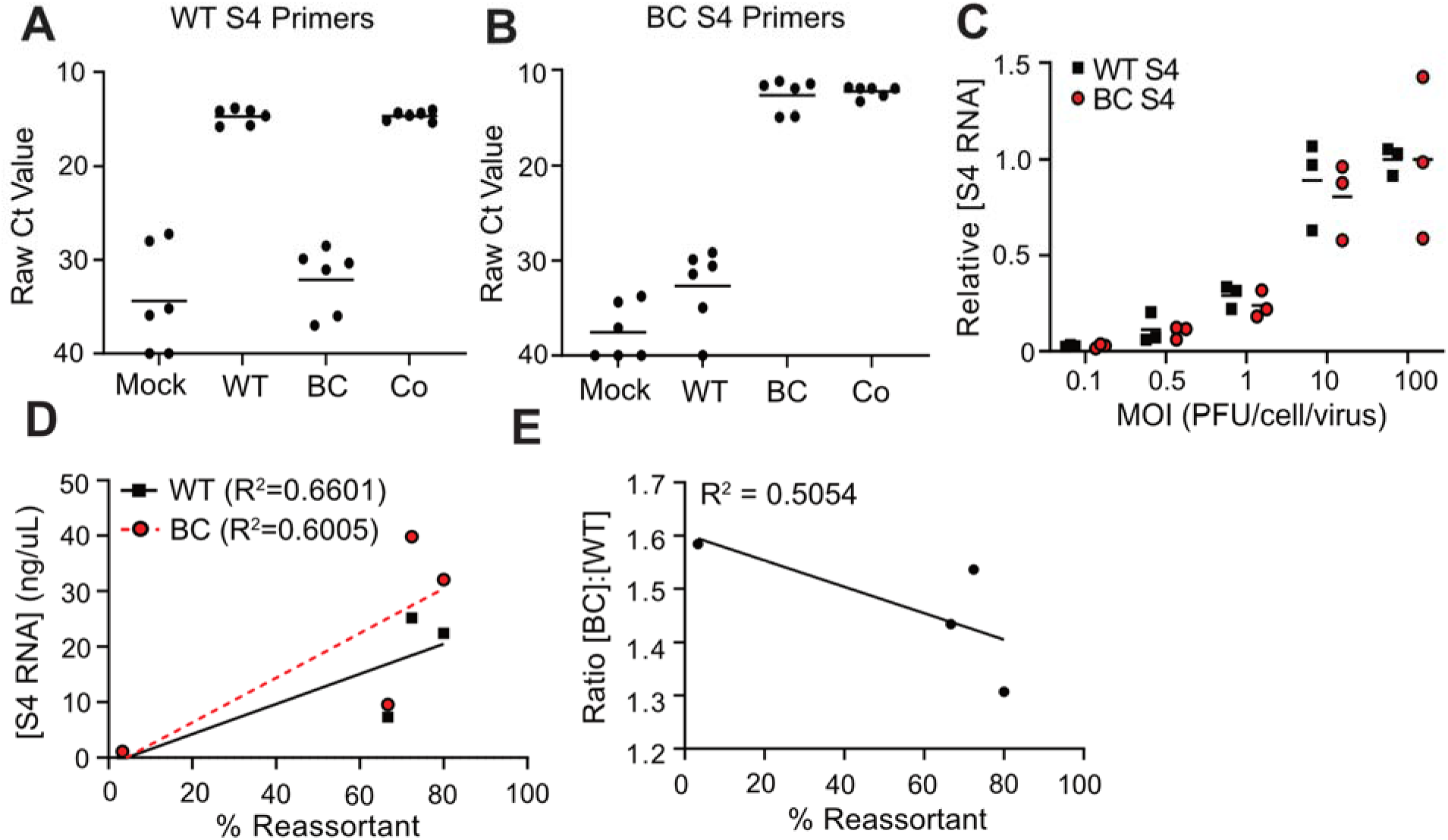
RNA abundance increases in concert with coinfection multiplicity. L cells were adsorbed with the indicated viruses at MOI = 10 PFU per cell per virus. RNA was extracted, cDNA was generated, and WT S4-specific primers (A) or BC S4-specific primers (B) were used to amplify cDNA. C_T_ values are shown for each infection condition. (C) L cells were coinfected at indicated multiplicities with WT and BC reovirus for 24 h before quantifying the concentration of WT and BC S4 RNA by RT-qPCR and normalizing based on MOI = 100 WT and BC RNA concentration. n = 3 pairs of independently plaque-purified clones (D-E) Simple linear regression analyses correlating the concentration of WT S4 RNA, BC S4 RNA (D), or the ratio of BC S4 RNA to WT S4 RNA (E) to reassortment frequency over a range of coinfection multiplicities.

To determine effects of RNA abundance of the coinfecting viruses on reassortment frequency, we quantified WT and BC S4 transcripts after coinfection of L cells over a range of MOIs. We found that RNA abundance for both viruses increased exponentially until MOI = 10, at which point the amount of RNA from both viruses stabilized (**Fig. 2C**). Linear regression analyses indicate a modest positive correlation between RNA abundance and reassortment frequencies observed during coinfection at increasing multiplicity (WT R^2^=0.6601; BC R^2^=0.6005) (**Fig. 2D-E**). The ratio of BC:WT RNA showed a weak negative correlation to reassortment frequency ([BC]:[WT] R^2^=0.5054), and the linear model does not appear to fit the data (**Fig. 2F**). These data suggest that the dramatic increase in RNA transcripts from both viruses provides ample opportunity for reassortment. Further, while RNA abundance increases in concert with reassortment frequency, it incompletely explains the data, suggesting that other factors likely contribute to coinfection reassortment outcomes.

### Reovirus reassortment frequency decreases with time delay to superinfection

In nature, viruses may infect a host or cell at different times, and primary infection may induce the organization of virus factories and host responses that can influence secondary infection, with unknown effects on reassortment. To assess the effect of infection timing on reassortment, we coinfected cells with WT and BC reoviruses asynchronously at a MOI of 10 PFU per cell per virus with a range of times from 0-16 h separating primary and secondary infection (**Fig. 3A**). We then quantified reassortment frequency among infectious viral progeny at 24 h post primary infection using HRM. We found that as the time delay between primary and secondary infection was extended, reassortment frequency declined (**Fig. 3B-C**). During coinfection, roughly two-thirds of progeny were reassortants, mirroring what was observed previously (**Figs. 1E and 3B**). However, with a 4 h delay to superinfection, reassortment frequency decreased from 65% to 43%. Reassortment frequency continued to decrease with increasing time delays to superinfection, such that at the latest time point tested, when superinfection occurred 16 h post-primary infection, only a single reassortant clone was isolated from a total of 30, yielding a reassortment frequency of 3% (**Fig. 3B-C**). As during coinfection, small segments were equally likely to reassort as large and medium segments. We predict three potential explanations for the observed decrease in reassortment frequency with increasing time to superinfection. First, since the superinfecting virus had less time to replicate than the primary infecting virus, viral RNA abundance could drive the reduction in reassortment frequency. This idea is supported by the observation that as the time delay to superinfection increased, there was a corresponding decrease in the proportion of total segments that were derived from the superinfecting BC virus.

**Figure 3.**
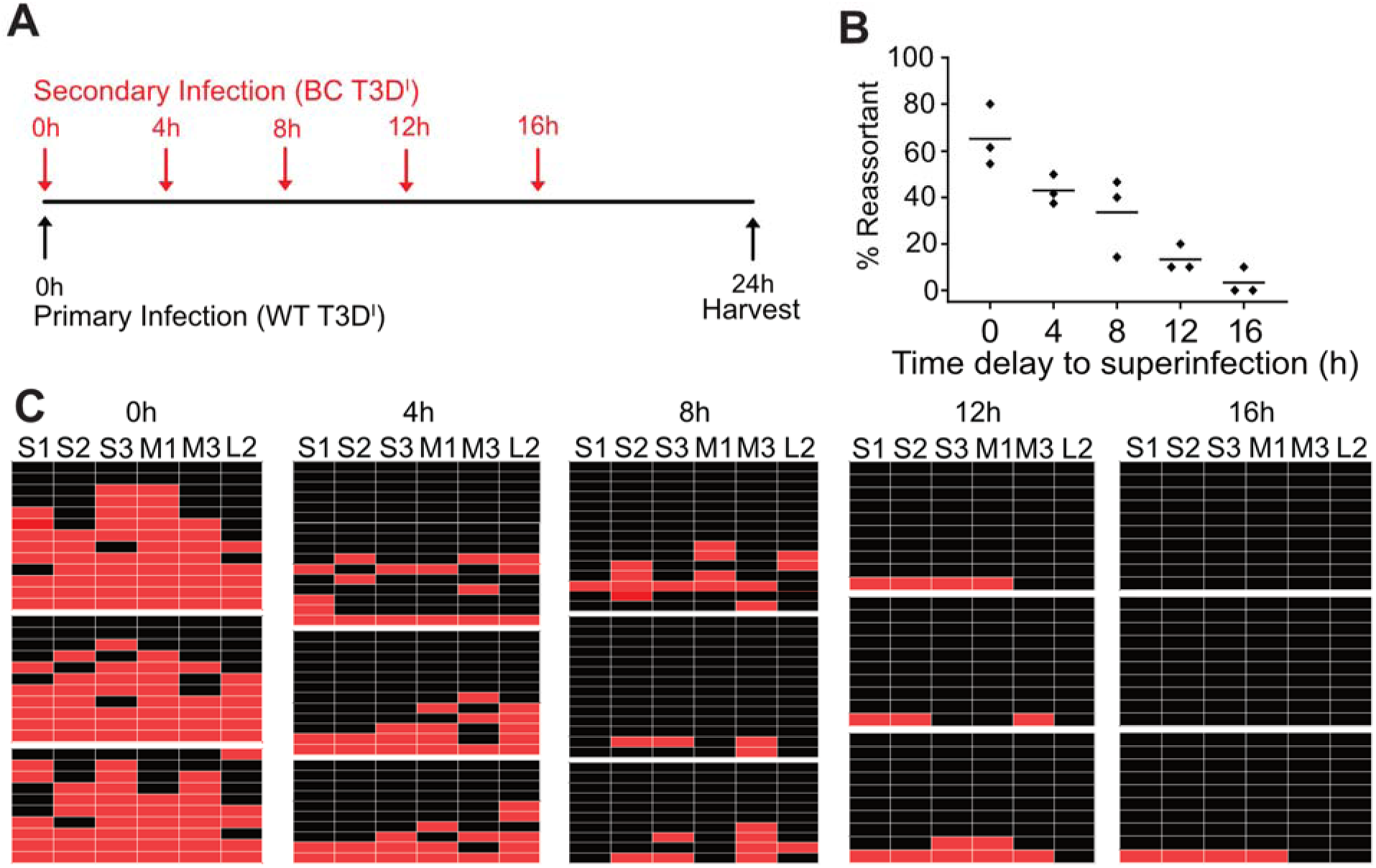
Reassortment frequency decreases with greater time delay to superinfection. (A) Schematic depicting the timing of coinfection and superinfection of L cells for superinfection time course. (B) L cells were adsorbed with WT prior to adsorption with BC at the indicated time post primary adsorption at MOI = 10 PFU per cell per virus. At 24 h post primary infection, reassortment frequency was quantified using HRM. (C) Honeycomb plots indicate genotypes of individual coinfection progeny viruses analyzed from the experiment in (B). Each row represents an independently isolated clone. Each column represents the indicated genome segment. Black indicates WT genome segments, and red indicates BC genome segments. n = 3 independent experiments with at least 10 progeny clones analyzed per experiment.

Next, it is possible that the observed decrease in reassortment frequency is a result of physical sequestration of +RNA within viral factories. Finally, it is possible that primary infection induces responses in the infected cell that prevent superinfection, such as the reduction of viral receptor expression or induction of interferon signaling.

### Viral RNA abundance correlates with reassortment frequency during superinfection

To quantify the abundance of viral RNA from each parental virus during superinfection we adsorbed L cells with WT reovirus and superinfected with BC reovirus at 0 (coinfection control), 4, 8, 12, or 16 h post primary infection. At 24 h post primary infection, we extracted total RNA and quantified S4 viral RNA abundance using RT-qPCR. We observed that while the abundance of RNA from the primary infecting WT virus remained stable or increased slightly over time, BC RNA abundance progressively declined, with the largest reduction in abundance coinciding with the longest time delays, consistent with reduced time for viral replication (**Fig. 4A**). Linear regression analysis revealed a weak negative correlation between the primary infecting virus RNA abundance and reassortment frequency (R^2^=0.5055) (**Fig. 4B**); that is, despite a slight increase in WT transcripts, reassortment events became increasingly rare over time.

**Figure 4.**
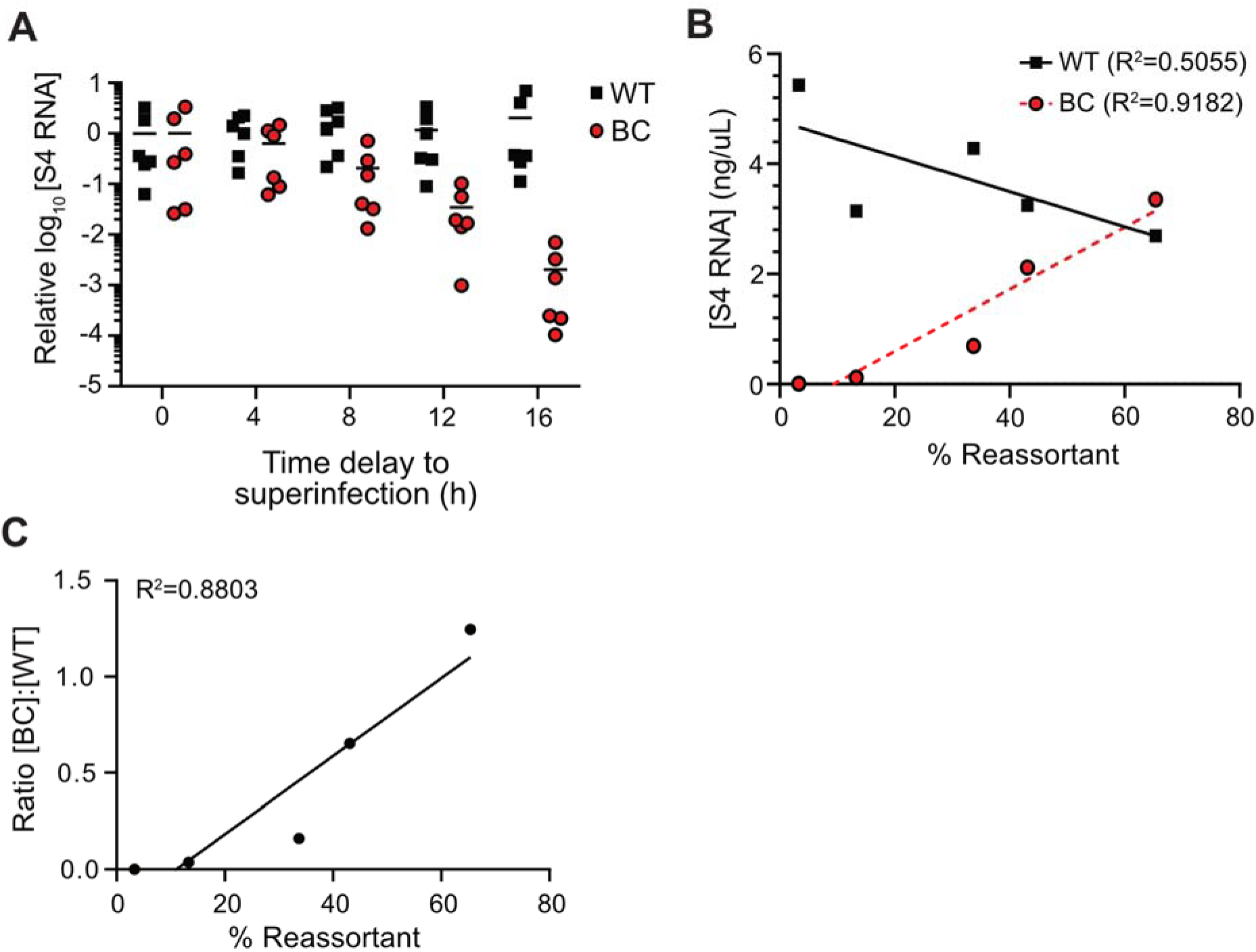
Superinfecting virus RNA abundance decreases with greater time to superinfection and correlates with reassortment frequency. (A) L cells were adsorbed with WT prior to adsorption with BC at the indicated time at MOI = 10 PFU per cell per virus. At 24 h p.i., WT and BC S4 RNA were quantified by RT-qPCR. (B) Simple linear regression analyses were used to correlate the concentration of WT S4 RNA or BC S4 RNA and (C) the ratio of BC:WT RNA to reassortment frequency at each superinfection time point. n = 3 pairs of plaque-purified clones from two independent experiments.

Linear regression analyses also showed that the abundance of BC RNA and the ratio of BC:WT RNA positively correlate in a linear manner with reovirus reassortment frequency (BC +RNA R^2^=0.9182; [BC]:[WT] R^2^=0.8803) (**Fig. 4C-D**). Therefore, the observed decrease in reassortment frequency can be explained by the substantial reduction in superinfecting virus RNA transcripts over time. However, there may be other contributing influences.

### Branched DNA FISH enables specific detection of WT and BC +RNA transcripts during coinfection

Coinfection of the host cell is required for two viruses to reassort genome segments. After observing that reovirus reassortment is non-random in the context of simultaneous coinfection, we sought to develop a tool with which we could quantify the percentage of cells that were infected with both WT and BC reoviruses during coinfection at increasing MOI. To detect infectivity by each virus, we designed differentially fluoresceinated branched DNA fluorescence *in situ* hybridization (bDNA FISH) probes specific to the WT and BC sequences of the S3, S4, and L1 genome segments. To determine if the probes were specific for their targets, we conducted high multiplicity single infections and coinfections with WT and BC, or a mock infection, fixed and permeabilized cells, stained nuclei with DAPI, and incubated all samples in the presence of both WT and BC probes to permit hybridization. Using immunofluorescence microscopy, we detected WT- or BC-specific probe binding in a substantial proportion of total cells only when the appropriate target virus was present, with low background binding to the non-target virus (**Fig. 5A-C)**. Although several fluorophores are conjugated to each bDNA FISH probe, since only a single probe can bind to a given viral +RNA molecule, it is unlikely that this method of detection provides single-molecule sensitivity (46). To compare the sensitivity of bDNA FISH for detecting total infectivity with that of antibody staining, we infected L cells with WT at increasing multiplicity and quantified infectivity using immunofluorescence microscopy after staining with either WT-specific bDNA FISH probes or with polyclonal reovirus antiserum and fluorescently-conjugated secondary antibodies. While the bDNA FISH approach could detect up to 80% +RNA-positive cells, it was less sensitive than immunostaining at most MOIs (**Fig. 5D**). Therefore, while bDNA FISH staining provides the benefit of detecting each virus individually during coinfection, its sensitivity likely is limited.

**Figure 5.**
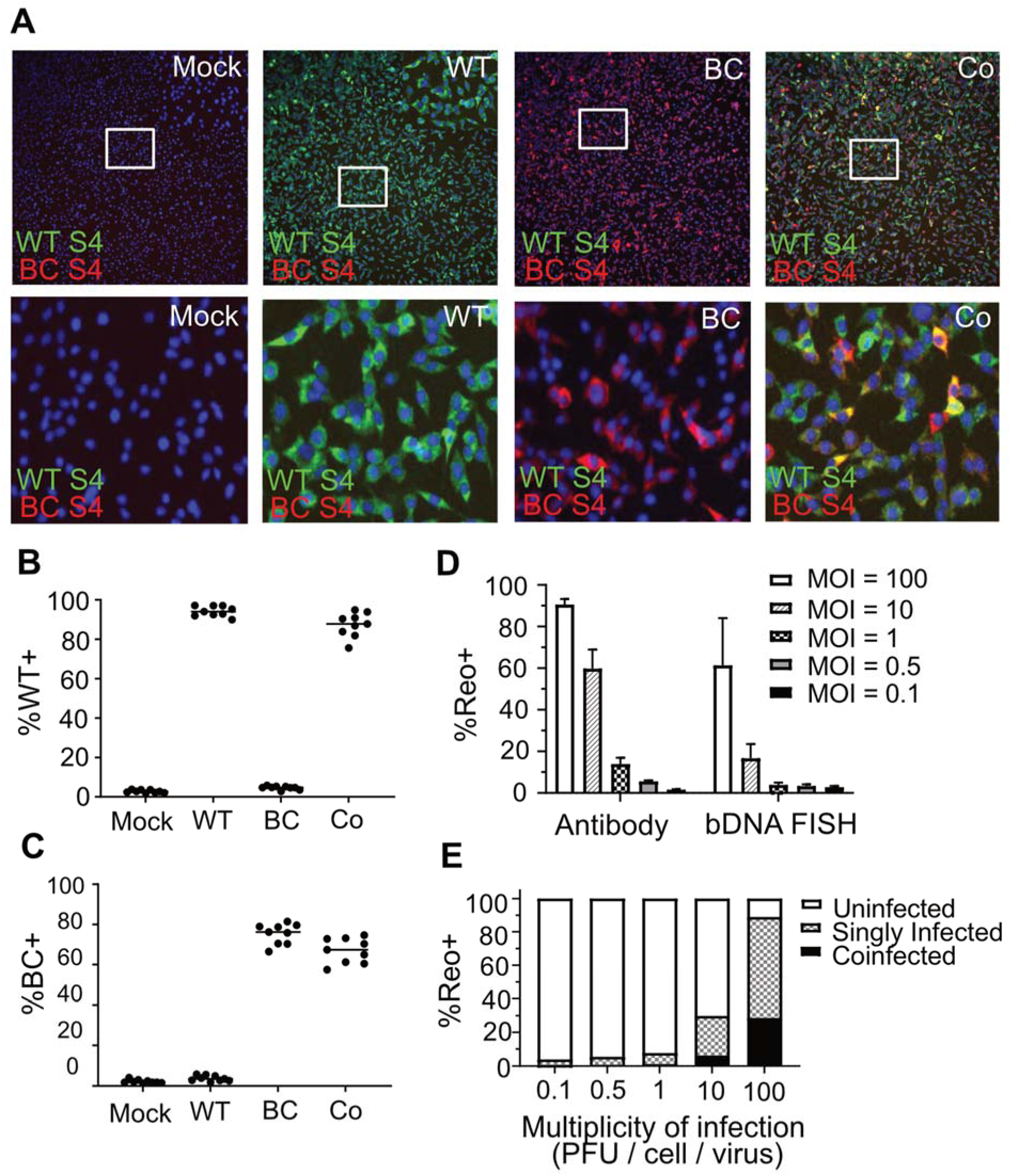
Branched DNA FISH enables specific detection of WT and BC +RNA during coinfection. L cells were adsorbed with medium (mock), WT only, BC only, or both WT and BC at a multiplicity of 10 PFU per cell per virus. Cells were fixed, stained for nuclei (blue), WT +RNA (green), and BC +RNA (red) using bDNA FISH probes, and visualized and quantified using an ImageXpress Micro high-content imaging system. Representative images are shown in (A). The percentage of total cells infected with WT (B) or BC (C) for each infection condition are indicated. n = 9 fields of view from one representative experiment. (D) L cells were adsorbed with the indicated MOI of WT reovirus and fixed and processed for imaging with the ImageXpress either using traditional immunostaining or branched DNA FISH workflow. The average percentage of total infected cells from four fields of view is shown. n = 3 plaque-purified clones. (E) L cells were coinfected at the indicated MOI with WT and BC reovirus for 24 h then fixed and processed for imaging with the ImageXpress using bDNA FISH workflow. The average percentage of uninfected, singly infected, and coinfected cells from four fields of view per clone are shown. n = 3 plaque-purified clones.

To determine the effect of MOI on codetection of viral +RNA, we adsorbed L cells with WT and BC reoviruses at increasing multiplicities and quantified the percentage of cells in which we could detect viral +RNA from a single virus or both viruses using bDNA FISH probes and immunofluorescence microscopy. The percent of cells in which we could detect +RNA from a single virus or from both viruses increased in concert with coinfection multiplicity, unlike the plateau observed for reassortment frequency at high MOIs (**Fig. 5E**). Even at the highest multiplicities assessed, 100% +RNA codetection was never achieved. While it is likely that the low sensitivity of the assay is in part responsible for this result, it may suggest that high levels of reassortment can occur under conditions of somewhat limited coinfection, or at least when limited amounts of both viral +RNAs are present. Considering that reassortment frequency began to plateau at a MOI of 1 PFU per cell per virus, while +RNA codetection continued to increase to a MOI of 100 PFU per cell per virus, factors other than +RNA codetection may impose barriers to entirely random reassortment.

### VFs are unlikely to influence reovirus reassortment frequency during superinfection

Reovirus establishes VFs within hours of entering host cells (26). Viral transcription is initiated in the cytoplasm but primarily occurs within VFs as early as 6 h p.i. (24). In addition, VFs act as sites of reovirus genome packaging and new particle assembly (24). Therefore, it is possible that newly synthesized +RNA transcripts from coinfecting viruses may be isolated within distinct VFs, posing a physical barrier to reassortment. However, VFs also display liquid-like properties whereby these structures undergo fusion and fission events (27), which could effectively promote reassortment events between viruses that are isolated to distinct VFs. To determine whether VFs introduce a physical barrier to co-localization of coinfecting viral +RNA during superinfection, and thereby reassortment, we assessed +RNA localization from coinfecting WT and BC reoviruses using bDNA FISH and confocal microscopy. WT virus was used as the superinfecting virus because the fluorophores conjugated to WT- specific probes were of a shorter wavelength, making it possible to detect superinfecting virus +RNA at late time points of superinfection. Specifically, we adsorbed L cells with BC reovirus and then superinfected with WT either at 0 (coinfection), 8, or 16 h post primary infection. We determined WT and BC +RNA localization by bDNA FISH at 24 h post primary infection using probes specific to the S3, S4, and L1 genome segments. At all time points of superinfection, +RNA from the superinfecting virus could be observed in the cytoplasm and within factories that were occupied by +RNA from the primary infecting virus (**Fig. 6A-C**). However, as the time delay to superinfection was increased, the percentage of factories within coinfected cells in which we detected +RNA from the superinfecting virus decreased (**Fig. 6D**). This decrease corresponded with a decrease in the percentage of VFs in which we detected +RNA from both the primary infecting and secondarily infecting viruses (**Fig. 6E**), and the ratio of VFs that were positive for WT +RNA relative to those that were positive for +RNA from both viruses remained constant at each time point (**Fig. 6F**). This finding suggests that the decrease in WT +RNA within VFs was responsible for the reduction in co-positive VFs. It was unclear, however, whether this reduction in the percentage of VFs containing +RNA from both viruses was due to superinfecting virus +RNA being excluded from existing VFs or was due to a reduction in superinfecting virus +RNA present within infected cells. To determine whether superinfecting virus +RNA was selectively excluded from VFs, we quantified the proportion of the total fluorescence intensity from superinfecting virus +RNA that was localized to VFs at each time point. We found that there were 22% and 27% reductions in sample means at 8 h and 16 h relative to 0 h, respectively, in the proportion of total WT superinfecting virus +RNA that localized to VFs (**Fig. 6G**). This represented a 7% reduction in superinfecting virus +RNA localized to VFs from 8h to 16h. However, there was an ∼10-fold decrease in reassortment frequency from 8 h to 16 h (8 h = 34% reassortant; 16 h = 3% reassortant) (**Fig. 3B**). Therefore, if superinfecting virus +RNA is excluded from VFs, it is only a small proportion of total +RNA and is unlikely to drive the observed reductions in reassortment frequency. The proportion of +RNA from the primary infecting virus localizing to VFs remained constant at each superinfection time point (**Fig. 6H**). Further analyses of +RNA within VFs revealed that superinfecting virus +RNA localized to progressively larger VFs as the time delay to superinfection increased (**Fig. 6I**). In contrast, +RNA from the primary infecting virus localized to VFs of roughly the same size at each superinfection time point (**Fig. 6J**). Additionally, the mean area of VFs containing +RNA from both viruses increased with greater time delay to superinfection (**Fig. 6K**). This finding provided additional evidence that large VFs do not exclude superinfecting virus +RNA, which suggests they do not preclude reassortment events. The mechanism through which superinfecting virus +RNA accesses mature VFs is open to further inquiry.

**Figure 6.**
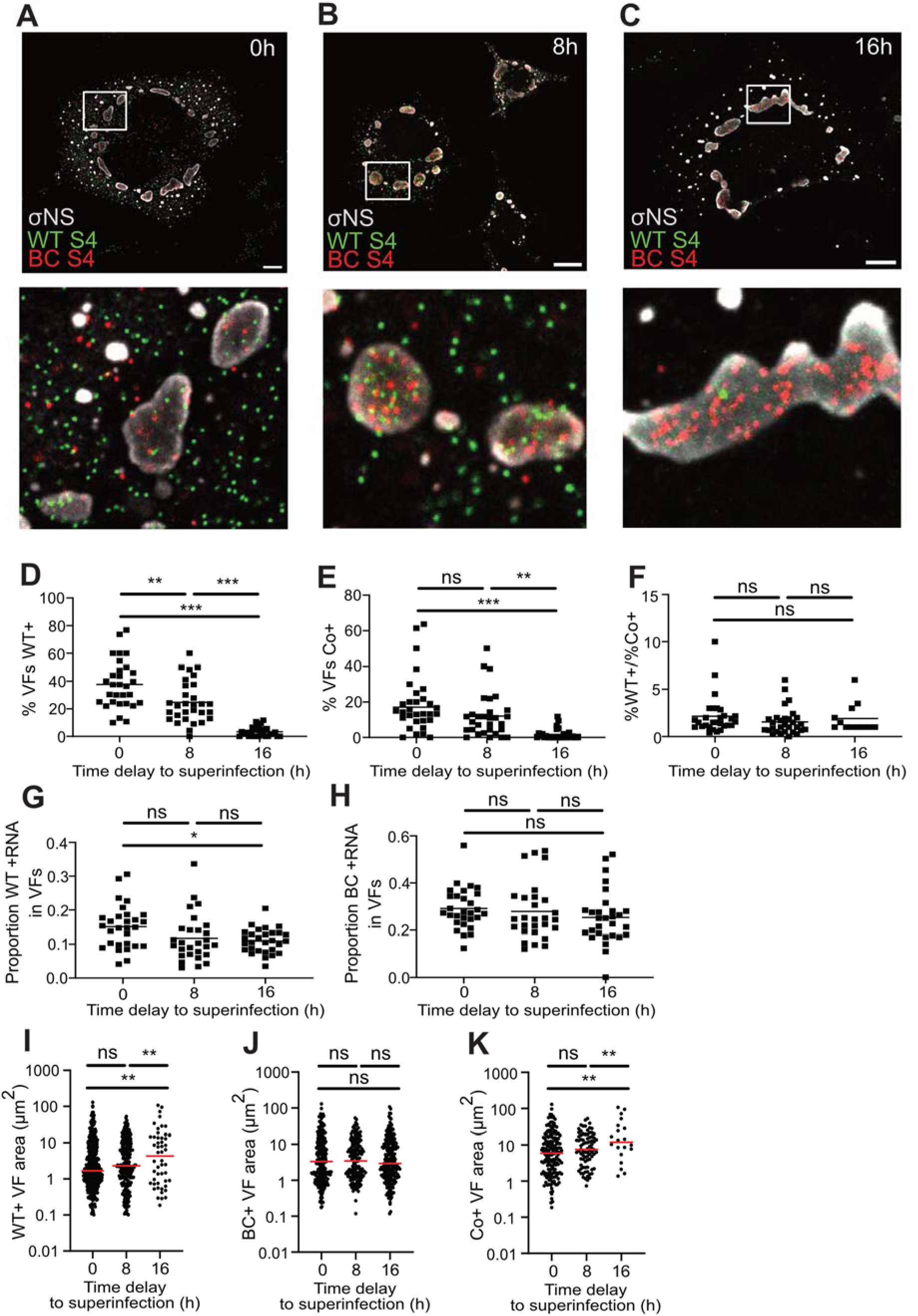
VFs do not exclude superinfecting virus +RNA. (A-C) L cells were coinfected with WT and BC reovirus simultaneously (A) or were infected at 0 h with BC and superinfected with WT 8 h (B) or 16 h (C) post primary infection. Cells were fixed and stained using bDNA FISH probes specific for S3, S4, and L1 +RNA of primary infecting BC (red) and superinfecting WT (green) and with antibodies against viral nonstructural protein σNS (gray) to define VFs. Cells were imaged using an LSM880 confocal microscope. Scale bar is 10 µm. (D-H) In 30 cells per condition, cells were segmented, and individual VFs were identified by thresholding based on σNS staining. WT and BC +RNA within all VFs and the cytoplasm were quantified using Fiji. The percentage of VFs that contain WT +RNA (D) or both WT and BC +RNA (E) is shown. The ratio of VFs that contain WT +RNA to those that contain both WT and BC +RNA is depicted in (F). The proportion of total fluorescence intensity from WT +RNA probes (G) and BC +RNA probes (H) that is localized to VFs at each superinfection time point is shown. (I-K) The average area of VFs positive for WT +RNA (I) BC +RNA (J), or both WT and BC +RNA (K) at the 0 h, 8 h, and 16 h superinfection time point is indicated. Significance determined by one-way ANOVA with Tukey’s multiple comparisons test. n = 30 cells per time point. * = p<0.05, ** = p<0.01, *** = p<0.001.

### T3D^I^ reovirus primary infection can limit superinfection

T3D reovirus induces host expression of type I and type III interferons in response to infection and is also sensitive to the effects of interferon (36, 37, 47, 48). Thus, we sought to determine whether type 3 reoviruses restrict reovirus superinfection. To first address this question, we adsorbed L cells with BC reovirus or medium alone (mock) at a MOI of 10 PFU per cell, coinfected or superinfected with WT reovirus at 0, 4, 8, or 16 h post primary infection, and quantified viral transcripts at 24 h p.i. Compared to the mock primary infection control, primary infection with BC reovirus had little to no effect on the abundance of viral transcripts generated by the superinfecting WT virus (**Fig. 7A**). However, maximal expression of IFNβ in response to reovirus infection does not occur until about 24 h p. i. (35). Therefore, we conducted an additional experiment in which we delayed the time to superinfection to 24 h post primary infection and allowed the secondary virus to replicate for 24 h. There was an ∼10-fold decrease in superinfecting virus transcript abundance following primary infection at MOI = 10 PFU/cell, relative to mock-infected controls (**Fig. 7B**). Thus, over the time course during which we quantified reassortment frequency (**Fig. 3C**), superinfection exclusion is unlikely to have driven the observed reduction in reassortment frequency. However, given a sufficient time delay to superinfection, primary infection with type 3 reovirus can restrict superinfecting virus replication, either through antiviral host responses or some other mechanism.

**Figure 7.**
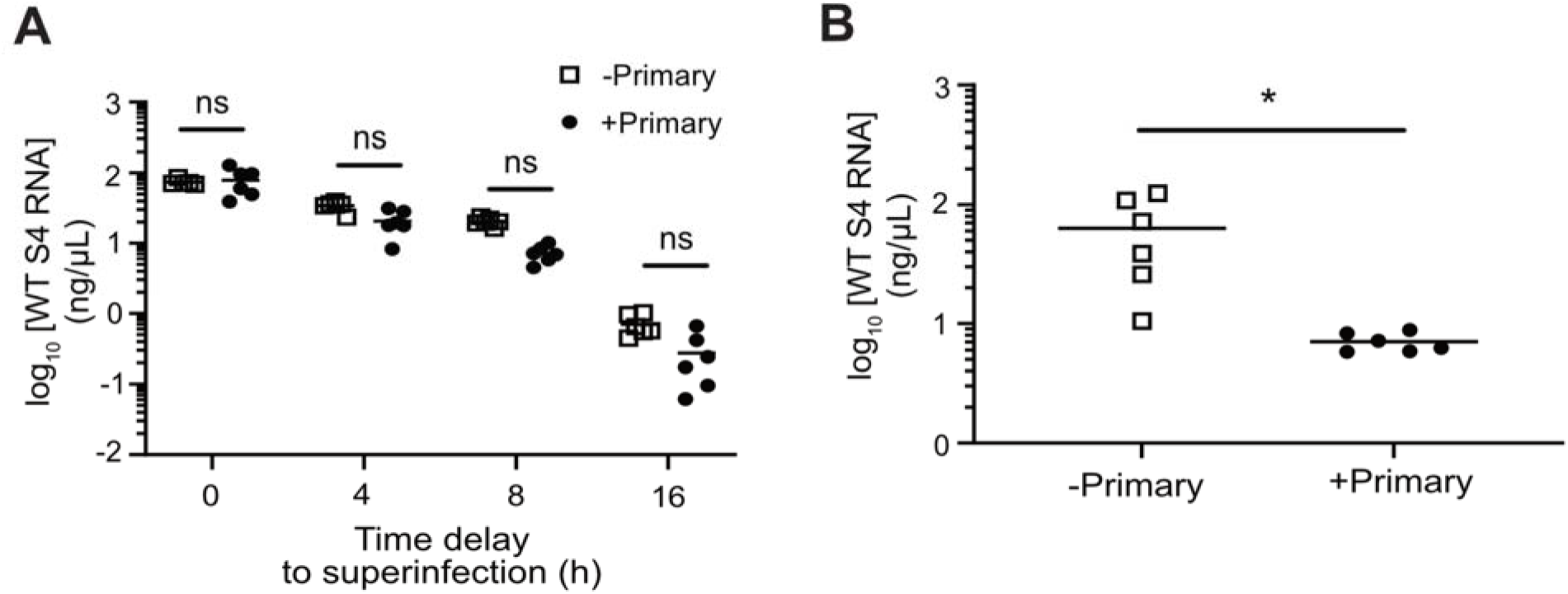
Superinfection is only inhibited by T3D^I^ reovirus with a 24 h primary infection. (A) L cells were adsorbed with medium (mock) or BC reovirus prior to adsorption with WT at the indicated time points. Superinfecting WT virus RNA concentration was quantified 24 h post primary infection by RT-qPCR. n = 3 plaque-purified clones from each of two independent experiments (B) L cells were adsorbed with medium (mock) or BC and incubated for 24 h prior to adsorption with WT. Superinfecting WT virus RNA concentration was quantified 48 h post primary infection by RT-qPCR. n = 3 plaque-purified clones from each of two independent experiments. Statistical significance was determined by two-way ANOVA with Sidak’s multiple comparisons test (A) and unpaired t-test (B) (* = p<0.05).

## DISCUSSION

In the present study, we show that type 3 reoviruses reassort genome segments efficiently during coinfection and superinfection. To achieve this, +RNA transcripts from superinfecting reovirus gain access to dense, cytoplasmic virus factories, facilitating generation of reassortant progeny. Additionally, superinfecting virus replication is not limited by antiviral responses initiated by type 3 reovirus within a single cycle of replication, though superinfection exclusion was observed and could influence reassortment with greater time delay to superinfection.

We observed that reassortment between wild-type and genetically-barcoded type 3 reoviruses is frequent during high multiplicity coinfection (**Fig. 1D**), consistent with published findings (28). RNA abundance failed to explain reassortment frequency during simultaneous coinfection (**Fig. 2D**), and the primary determinant of reassortment frequency during coinfection remains unclear. Coinfection of a cell is required for reassortment to occur. Thus, coinfection frequency is likely an important driver of reassortment frequency. However, quantitation of coinfection frequency with genetically-similar viruses remains technically difficult (28). Our bDNA probes are insufficiently sensitive to accurately quantify coinfection for WT and BC viruses, which renders correlations between coinfection frequency and reassortment frequency dubious. We also observed that reassortment decreases in frequency with increasing time delay to superinfection. We proposed that this reduction in reassortment frequency could be explained by the fact that transcripts from the superinfecting virus are less abundant under the infection conditions or could be the product of some other limiting intrinsic influence, including transcript compartmentalization in VFs or superinfection exclusion. Under the conditions tested, in which superinfection occurs within a single cycle of replication by the primary virus (<16 h), we found that the decrease in superinfecting virus transcripts strongly correlated with reassortment frequency (**Fig. 4B**), offering a potential explanation for why reassortment becomes less frequent with greater time delay to superinfection.

Nascent reovirus +RNA transcripts are generated after reovirus core particles are deposited into the cytoplasm (49–51), and reovirus transcription is primarily localized to VFs as early as 6 h p.i. (24). Reovirus VFs form early in infection following transcription and translation of nonstructural proteins µNS and σNS, which associate with and recruit viral RNA and transcriptionally-active core particles (26, 39, 52, 53). Thus, we initially anticipated that most viral +RNA would be localized to VFs and that +RNA from coinfecting viruses would quickly compartmentalize into distinct VFs, posing a barrier to reassortment. In contrast, we observed that reovirus +RNA localizes both to VFs and the cytoplasm and that +RNA from coinfecting and superinfecting viruses frequently co-occupied VFs (**Fig. 6A-C)**. Since initiating our study, others have also observed that reovirus +RNA localizes to both VFs and the cytoplasm (25). The mechanism through which +RNA from coinfecting viruses acquires access to the same VF remains unclear.

However, reovirus core particles can be recruited to factory-like structures formed by µNS (52), and the nonstructural protein σNS recruits viral RNA to VFs (25). Thus, µNS and σNS may indiscriminately recruit cores and cytoplasmic pools of viral +RNA from distinct viruses to the same VF, providing opportunity for reassortment. It is also possible that VF fusion events facilitate co-occupation of VFs by +RNA from coinfecting viruses. Rotavirus viroplasms display properties of liquid condensates (54), and reovirus VFs similarly undergo fusion and fission events which rely on stabilized microtubules (27). However, disruption of VFs with microtubule-depolymerizing drugs does not limit reassortment frequency in the context of simultaneous coinfection (28), suggesting that VF fusion is not required for +RNA from coinfecting viruses to occupy the same VF. Importantly, we also found that superinfecting virus +RNA was not strictly localized to small VFs but was instead localized to VFs of a range of sizes (**Fig. 6I**). This observation suggests either that superinfecting virus does not establish distinct VFs, or that superinfecting virus establishes distinct VFs that quickly fuse with existing VFs.

Further, the mean area of a VF occupied by +RNA from the superinfecting virus increased at later times of addition (**Fig. 6I**). The simplest explanation for this finding is that it is the result of omission bias; when there is less +RNA in the cell, it is more likely that +RNA will be detected in VFs that occupy a larger area. Given that the bDNA probes used were not sufficiently sensitive to detect all +RNA transcripts, probe sensitivity almost certainly has some effect. However, it is also possible that larger VFs, occupying a greater surface area, are more likely to be near incoming cores and recently transcribed +RNA, and as such are more likely to recruit superinfecting virus. Finally, rotavirus viroplasms lose their liquid-like properties later in infection, coinciding with phosphorylation of NSP5 (54). If reovirus VFs behave similarly, this might suggest that established VFs are less likely to undergo fusion events, and recruitment of superinfecting virus +RNA to large VFs likely is not due to fusion of small VFs with larger VFs.

Our findings suggest that superinfection exclusion is unlikely to influence reassortment frequency during the time course tested but may have an effect when secondary infection is introduced with greater time delays. Previous explorations of *Reoviridae* virus superinfection exclusion have yielded mixed results. Prior work found that reassortment still occurs between coinfecting viruses with a time delay to superinfection up to 24 h, indicating that superinfection is not completely precluded by reovirus primary infection (34). A similar observation has also been made for rotavirus (55). In contrast, superinfection exclusion studies involving bluetongue virus indicate that primary infection restricts superinfection both *in vitro* and *in vivo* and that it does so as early as 4 h post-primary infection *in vitro*, at least upon secondary infection with extracellular vesicle-associated virus (56–58). The mechanism of reovirus superinfection exclusion is unclear. Known mechanisms of superinfection exclusion include antiviral host responses (32, 33), inhibition of viral entry through various mechanisms (17, 31, 59, 60), and competition for host resources (29). Given that type 3 reoviruses are sensitive to type 1 interferon (36), antiviral responses seem a likely mechanism for superinfection exclusion. Mutations in µ2 which cause T1L reovirus to induce interferon responses similar to T3D (61, 62) did not yield significant changes in reassortment frequency during simultaneous coinfection (28). Thus, preexisting antiviral responses may be required to limit viral replication to such an extent that reassortment is also inhibited.

Reovirus could also limit superinfection through other mechanisms. In the present study, exclusion was observed during high multiplicity superinfection and after a complete replication cycle. Given the abundance of new virus being generated under these conditions, it is possible that finite host resources are a limiting factor in the ability of superinfecting viruses to replicate. The timing of superinfection exclusion may also be telling. Superinfection was inhibited 24 h after primary infection and not with shorter time delays. Apoptosis limits type 3 reovirus replication in the intestine (63). However, only a small percentage of cells undergo apoptosis and necroptosis by 24 h following type 3 reovirus infection (64–66). Thus, cell death is unlikely to drive the observed exclusion. Bluetongue virus superinfection exclusion is overcome by free virus particles, but not extracellular vesicle-associated virus, and as such superinfection may be restricted at the step of virus entry (56). However, given that the timing of superinfection exclusion onset differs between bluetongue virus and reovirus, the mechanism of superinfection exclusion for these viruses may not be shared.

What these findings mean for reassortment in nature is an open question. Reassortment events are considered disadvantageous to the virus in most instances, as reassortment can disrupt conserved RNA and protein interactions that are essential for virus replication (67, 68), and reassortant influenza viruses have been shown to have fewer descendants than non-reassortants (69). However, reassortant viruses are frequently identified in nature, and reassortment events are common in the evolutionary history of many viruses (9, 15, 69). In the current study, we analyzed reassortment in the absence of segment mismatch. These findings reveal distinct sets of influences from those at play when coinfecting virus sequences are highly divergent and may be more applicable to intrapopulation or intrasubtype reassortment, which would involve highly genetically similar viruses. Influenza A virus reassortment has been shown to be under distance-dependent negative selection – that is, reassortment is more detrimental to progeny fitness when parent viruses are genetically dissimilar (69). In the context of genetically-similar viruses, we detected few substantial restrictions to reassortment during coinfection and superinfection *in vitro*. Our work contributes to a growing literature suggesting that reassortment is tolerated and efficient for viruses that are highly similar (28, 44). Whether reovirus has evolved to promote reassortment, or whether reassortment is simply tolerated as a by-product of the reovirus replication strategy, remains to be determined.

## ACKNOWLEDGEMENTS

We thank the staff at the Vanderbilt Cell Imaging Shared Resource for assistance with confocal microscopy image acquisition and analysis. We also thank the Vanderbilt High-Throughput Screening group for assistance with infectivity and coinfectivity assays using the ImageXpress platform. We appreciate the efforts of Dr. Terence Dermody, Dr. Alexa Roth, and Alejandra Flores for critical reading of the manuscript.

This work was supported by the National Institutes of Health (1R01AI155646 to K.M.O.), by a scholarship from the Vanderbilt Digestive Disease Research Consortium (to K.M.O), and by the Vanderbilt Institute for Clinical and Translational Research (VR53855 to T.W.T.) which is supported by the National Center for Advancing Translational Sciences (CTSA Award No. UL1 TR002243). The contents of this publication are solely the responsibility of the authors and do not necessarily represent the views of the National Center for Advancing Translational Sciences and the National Institutes of Health.

